# Acid sensing ion channel 1a is a key mediator of cardiac ischemia-reperfusion injury

**DOI:** 10.1101/869826

**Authors:** Meredith A. Redd, Sarah E. Scheuer, Natalie J. Saez, Ling Gao, Mark Hicks, Jeanette E. Villanueva, Melissa E. Reichelt, Jason N. Peart, Louise E. See Hoe, Han S. Chiu, Xiaoli Chen, Yuliangzi Sun, Jacky Y. Suen, Robert J Hatch, Ben Rollo, Mubarak A.H. Alzubaidi, Snezana Maljevic, Greg A. Quaife-Ryan, Walter G. Thomas, James E. Hudson, Enzo R. Porrello, Gabriel Cuellar-Partida, John F. Fraser, Steven Petrou, Glenn F. King, Peter S. Macdonald, Nathan J. Palpant

**Affiliations:** Institute for Molecular Bioscience, The University of Queensland, St. Lucia, Australia; Victor Chang Cardiac Research Institute, Sydney, Australia; Cardiopulmonary Transplant Unit, St. Vincent’s Hospital, Sydney, Australia; Faculty of Medicine, University of New South Wales, Sydney, Australia; Department of Pharmacology, St. Vincent’s Hospital, Sydney, Australia; School of Biomedical Sciences, The University of Queensland, St. Lucia, Australia; School of Medical Science, Griffith University, Southport, Australia; Faculty of Medicine, The University of Queensland, Brisbane, Australia; Critical Care Research Group, The Prince Charles Hospital and The University of Queensland, Brisbane, Australia; Florey Institute of Neuroscience and Mental Health, University of Melbourne, Melbourne, Australia; QIMR Berghofer Medical Research Institute, Brisbane, Australia; Murdoch Children’s Research Institute, The Royal Children’s Hospital, Melbourne, Australia; Department of Physiology, School of Biomedical Sciences, The University of Melbourne, Parkville, Australia; The University of Queensland Diamantina Institute, Faculty of Medicine and Translational Research Institute, Woolloongabba, Australia

**Author notes:** These authors contributed equally (Meredith A. Redd, Sarah E. Scheuer). Corresponding authors, Addresses for correspondence: Dr. Nathan J. Palpant, Institute for Molecular Bioscience, 306 Carmody Road, Building 80 (Level 4 North), St. Lucia, Queensland, Australia, 4072; E, Professor Glenn F. King, Institute for Molecular Bioscience, 306 Carmody Road, Building 80 (Level 2 North), St. Lucia, Queensland, Australia, 4072; E, Professor Peter Macdonald, Cardiopulmonary Transplant Unit, St. Vincent’s Hospital Darlinghurst, New South Wales, Australia; E.

## Abstract

The proton-gated acid-sensing ion channel 1a (ASIC1a) is implicated in the injury response to cerebral ischemia but little is known about its role in cardiac ischemia. We provide genetic evidence that ASIC1a is involved in myocardial ischemia-reperfusion injury (IRI) and show that pharmacological inhibition of ASIC1a yields robust cardioprotection in rodent and human models of cardiac ischemia, resulting in improved post-IRI cardiac viability and function. Consistent with a key role for ASIC1a in cardiac ischemia, we show that polymorphisms in the ASIC1 genetic locus are strongly associated with myocardial infarction. Collectively, our data provide compelling evidence that ASIC1a is a key target for cardioprotective drugs to reduce the burden of disease associated with myocardial ischemia.

## Introduction

Conditions caused by obstruction of blood flow to the heart, such as myocardial infarction (MI), are the most common emergency manifestation of cardiovascular disease^1^. Although acute reperfusion therapies have improved patient outcomes, mortality remains high^2^ and MI is one of the largest attributable risks for heart failure (HF)^3^. Globally, 1 in 5 people develop HF, with annual healthcare costs of $108B^4,5^. Heart transplantation remains the most effective treatment option for HF^6,7^, but 75% of potential donor hearts are discarded, many due to sensitivity of the donor heart to ischemic injury^8^. Myocardial sensitivity to ischemia-reperfusion injury (IRI) therefore remains a primary point of vulnerability underlying cardiovascular disease, which is the leading cause of morbidity and mortality worldwide^9^. Despite decades of preclinical therapeutic development, there are currently no drugs that block the acute injury response to cardiac ischemia^10^.

Myocardial IRI is a complex pathophysiological process that underlies the cardiac injury sustained during cardiac surgery, heart transplant, MI, and cardiac arrest. During myocardial ischemia, reduced oxygen availability causes a shift from fatty acid metabolism to anaerobic glycolysis^11,12^. The resulting lactic acidosis causes the extracellular pH to fall to as low as 6.0– 6.5^13,14^. In patients suffering from acute MI, the severity of acidosis strongly correlates with patient mortality, with pH < 7.35 associated with > 60% mortality^15^. Ongoing acidosis results in the activation of several transmembrane ion channels and receptors, including the sodium-hydrogen exchanger (NHE), glutamate-gated NMDA receptors, and TRP family of ion channels, each of which are thought to contribute to calcium overload, metabolic dysfunction, and eventually cell death^16–18^. In line with this, the Expedition Trial showed significant protection against peri-operative MI with the NHE inhibitor, cariporide, in patients undergoing high-risk coronary artery bypass surgery^19^. While this trial failed due to cerebrovascular complications, it demonstrated that pharmacological conditioning has the capacity to reduce the injury response to myocardial IRI.

Acid sensing ion channels (ASICs) are voltage-independent proton-gated cation channels of the degenerin/epithelial sodium channel superfamily^20^. There are six ASIC isoforms derived from four genes (*ACCN1–4*), which assemble into homotrimeric or heterotrimeric channels. The pH sensitivity and kinetics of ASICs are determined by their subunit composition^21,22^. ASICs are involved in extracellular acidification-induced calcium overload in neurons and cardiomyocytes^23–25^. ASIC1a, a splice variant of the *ACCN1* gene, is the most pH-sensitive ASIC channel. Activation begins at pH ≤ 7, with half-maximal activation at pH 6.6^26,27^, and ASIC1a currents are potentiated by numerous metabolic events that occur during ischemia including membrane stretch and increased levels of extracellular lactate, pyruvate, and arachidonic acid^28,29^. ASIC1a plays a critical role in mediating neuronal death after cerebral ischemia, and inhibition of ASIC1a through genetic ablation or pharmacological blockade is neuroprotective in rodent models of ischemic stroke^30–32^.

In this study, we used genetic and pharmacological approaches to demonstrate, for the first time, that ASIC1a is a critical mediator of the injury response to myocardial IRI. We show that venom-derived inhibitors of ASIC1a provide robust cardioprotection across multiple species and model systems from the cellular to whole-organ level, including models of IRI and donor heart preservation. Taken together, our data provide compelling evidence that ASIC1a is an important target for cardioprotective drugs, with potential application in the treatment of MI, cardiac arrest, and the preservation of donor hearts.

## Results

### Genetic ablation of ASIC1a improves functional recovery following cardiac IRI

We assessed ASIC isoform expression in the adult mouse heart from transcriptomic data of sorted cardiac cell populations after sham and myocardial infarction (MI)^33^. In sham hearts, *ASIC1* expression was highest in cardiomyocytes, while endothelial cells and fibroblasts expressed both *ASIC1* and *ASIC3*. In all three cell types, *ASIC4* and *ASIC2* had low and undetectable expression, respectively (**Fig. 1a**). *ASIC1* was not differentially expressed between sham and post-MI conditions in cardiomyocytes or endothelial cells (**Supplementary Fig. 1a**). *ASIC1* encodes two isoforms from the same genetic locus, ASIC1a and ASIC1b, that are genetically conserved among bilateria. As opposed to ASIC1b, which is primarily involved in nociception^34^, we focused on ASIC1a due to its known role in mediating injury responses resulting from ischemia in the brain^22^. We subsequently tested whether ASIC1a governs the response to cardiac ischemic injury.

**Figure 1.**
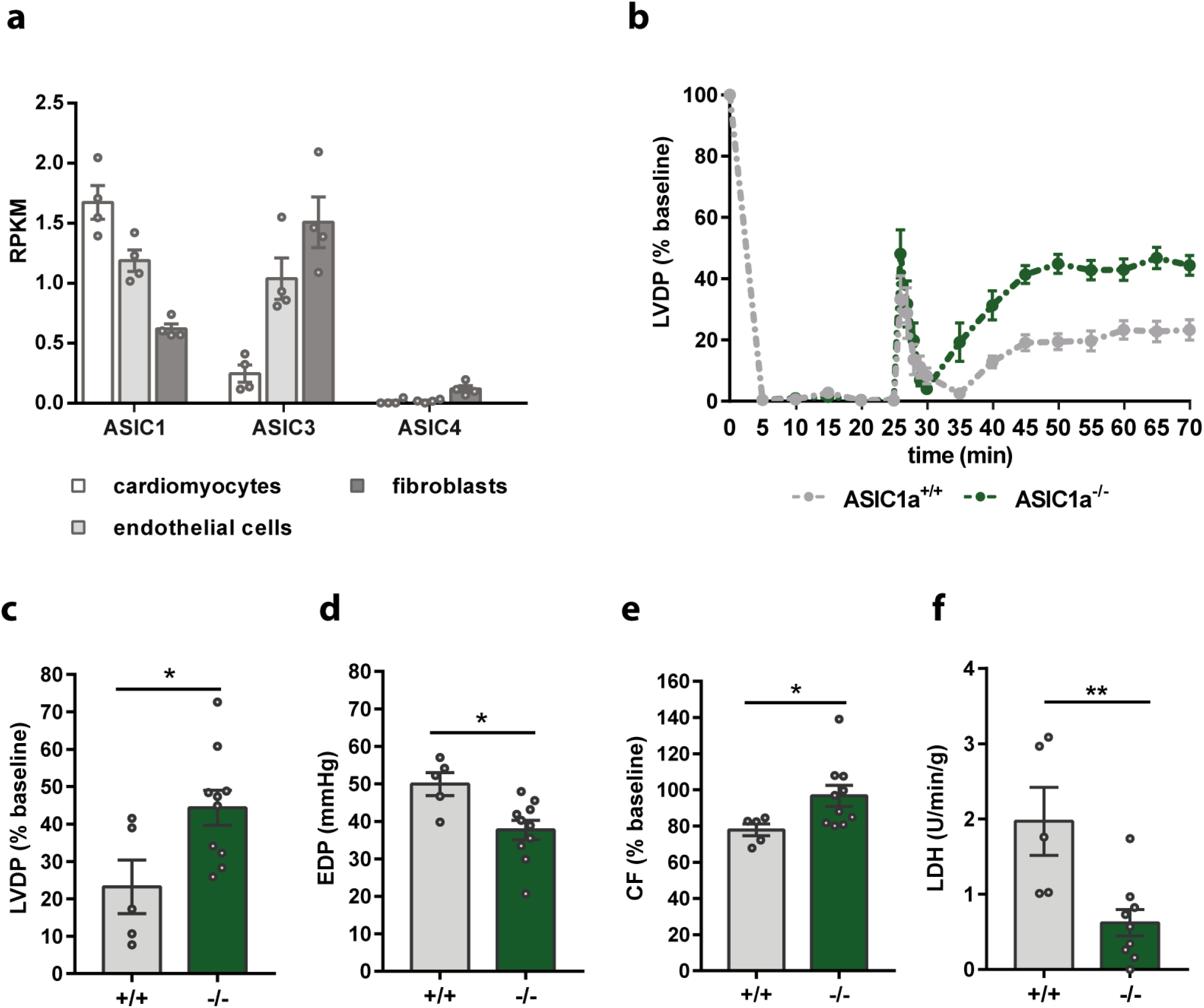
Genetic knockout of ASIC1a protects mouse hearts from *ex vivo* IRI. (**a**) mRNA expression (reads per kilobase million, RPKM) analysis of sorted cardiac cell populations from P56 adult mouse hearts (data extracted from Ref.^33^). ASIC2 was not detected. (**b-f**) Langendorff-perfused hearts from ASIC1a KO (ASIC1a^-/-^, *n* = 10) and WT (ASIC1a^+/+^, *n* = 5) mice were subjected to 25 min of global ischemia followed by 45 min of reperfusion. (**b**) LVDP, expressed as % of pre-ischemia baseline, over time. (**c-e**) Functional parameters measured or calculated after 45 min of reperfusion. (**c**) LVDP, percent baseline (p = 0.025). (**d**) EDP, p = 0.013. (**e**) CF, percent baseline (p = 0.049). (**f**) Cell death after 2 min of reperfusion (units/mL of LDH normalized to reperfusion flow rate and heart weight, U/min/g, p = 0.006). For LVDP and CF, baseline values were obtained immediately prior to ischemia. All data are expressed as mean ± SEM. Statistical significance was determined using two-tailed unpaired student’s *t*-test (*p < 0.05; **p < 0.01).

To determine whether ASIC1a plays a functional role during cardiac ischemia, we assessed IRI tolerance of Langendorff-perfused isolated hearts from ASIC1a-specific knockout (KO) mice. To generate the ASIC1a KO mouse strain, we used CRISPR editing to target the *ACCN1* locus. Specificity of the knockout was confirmed using qRT-PCR that showed only *ASIC1a* but not *ASIC1b* mRNA was reduced in brain tissue from knockout mice (**Supplementary Fig. 2**). Baseline function and heart rate in ASIC1a KO (ASIC1a^-/-^) isolated hearts were comparable to those measured in wildtype (WT) control hearts (ASIC1a^+/+^) (**Table 1**). To assess tolerance to IRI, hearts were subjected to 25 min of global normothermic zero-flow ischemia, followed by 45 min of aerobic reperfusion. WT and ASIC1a KO hearts showed similar initial responses to ischemia with comparable levels of ventricular contracture (**Supplementary Fig. 1b**). During reperfusion, both groups demonstrated gradual recovery of contractile function over time, with markedly improved functional recovery in ASIC1a KO hearts (**Fig. 1b**). By the end of the reperfusion period, ASIC1a KO hearts had significantly improved systolic and diastolic function with higher left ventricular developed pressure (LVDP) (44 ± 5% of baseline) and lower end diastolic pressure (EDP) (38 ± 3 mmHg) compared to WT hearts (LVDP: 23 ± 7% of baseline; EDP: 50 ± 3 mmHg); **Fig. 1c,d**). ASIC1a KO hearts also had improved recovery of coronary flow (CF) rates by the end of reperfusion (KO: 97 ± 6%; WT: 78 ± 3%, percent baseline) (**Fig. 1e** and **Supplementary Fig. 1c,d**). To assess cell death, lactate dehydrogenase (LDH) efflux was measured in perfusate samples collected during reperfusion. LDH efflux from ASIC1a KO hearts was significantly lower compared to WT hearts after 2 min of reperfusion (**Fig. 1f**), with a similar trend at the end of reperfusion (**Supplementary Fig. 1d**). Our data indicate that ASIC1a does not play a role in maintaining functional homeostasis, but it is a critical mediator of the organ response to myocardial IRI.

**Table 1.** Baseline functional parameters for Langendorff-perfused hearts from ASIC1a^+/+^ and ASIC1a^-/-^ mice

| Genotype | Heart Rate, bpm | LVDP, mmHg | EDP, mmHg | +dP/dt (mmHg/s) | -dP/dt (mmHg/s) | CF (mL/min/g) |
| --- | --- | --- | --- | --- | --- | --- |
| Wildtype (ASIC1a<sup>+/+</sup>) | 353 ± 23 | 105 ± 4 | 6.8 ± 0.7 | 3777 ± 139 | -2592 ± 144 | 35 ± 5 |
| KO (ASIC1a<sup>-/-</sup>) | 339 ± 17 | 101 ± 4 | 5.1 ± 0.7 | 3647 ± 236 | -2702 ± 148 | 26 ± 3 |
| *p-value* | 0.67 | 0.54 | 0.13 | 0.66 | 0.64 | 0.25 |

All pre-ischemia functional parameters were measured after more than 30 min of stabilization, except for heart rate, which was measured after 15 min of stabilization, prior to ventricular pacing. All values are expressed as mean ± SEM.

### ASIC1a inhibitors protect mouse hearts against IRI

We sought to determine whether pharmacological inhibition of ASIC1a is cardioprotective during an acute cardiac ischemic insult. Despite significant investment and preclinical testing, drug development pipelines have failed to identify effective small molecule inhibitors of ASIC1a due to lack of specificity, potency, or functional efficacy^35–37^. We therefore took advantage of two venom-derived inhibitors of ASIC1a, including Hi1a, a 76-residue double-knot peptide that we discovered from the venom of an Australian funnel-web spider (**Fig. 2a**). Hi1a is the most potent and selective inhibitor of ASIC1a identified to date, with an IC_50_ of 500 pM^32^. PcTx1, a single-knot peptide from the venom of a tarantula, is also a potent and selective inhibitor of ASIC1a (IC_50_ ~1 nM)^30^. Although the two peptides are closely related (Hi1a is comprised of two PcTx1-like domains joined by a short linker; **Fig. 2b,c**), they have distinct inhibitory modes of action. Hi1a inhibits ASIC1a activation whereas PcTx1 promotes and stabilizes a desensitized state of the channel^32^. We also utilized an analogue of PcTx1 that contains mutations of two pharmacophore residues (R27A/V32A), which dramatically reduces its inhibitory effect on ASIC1a^30^.

**Figure 2.**
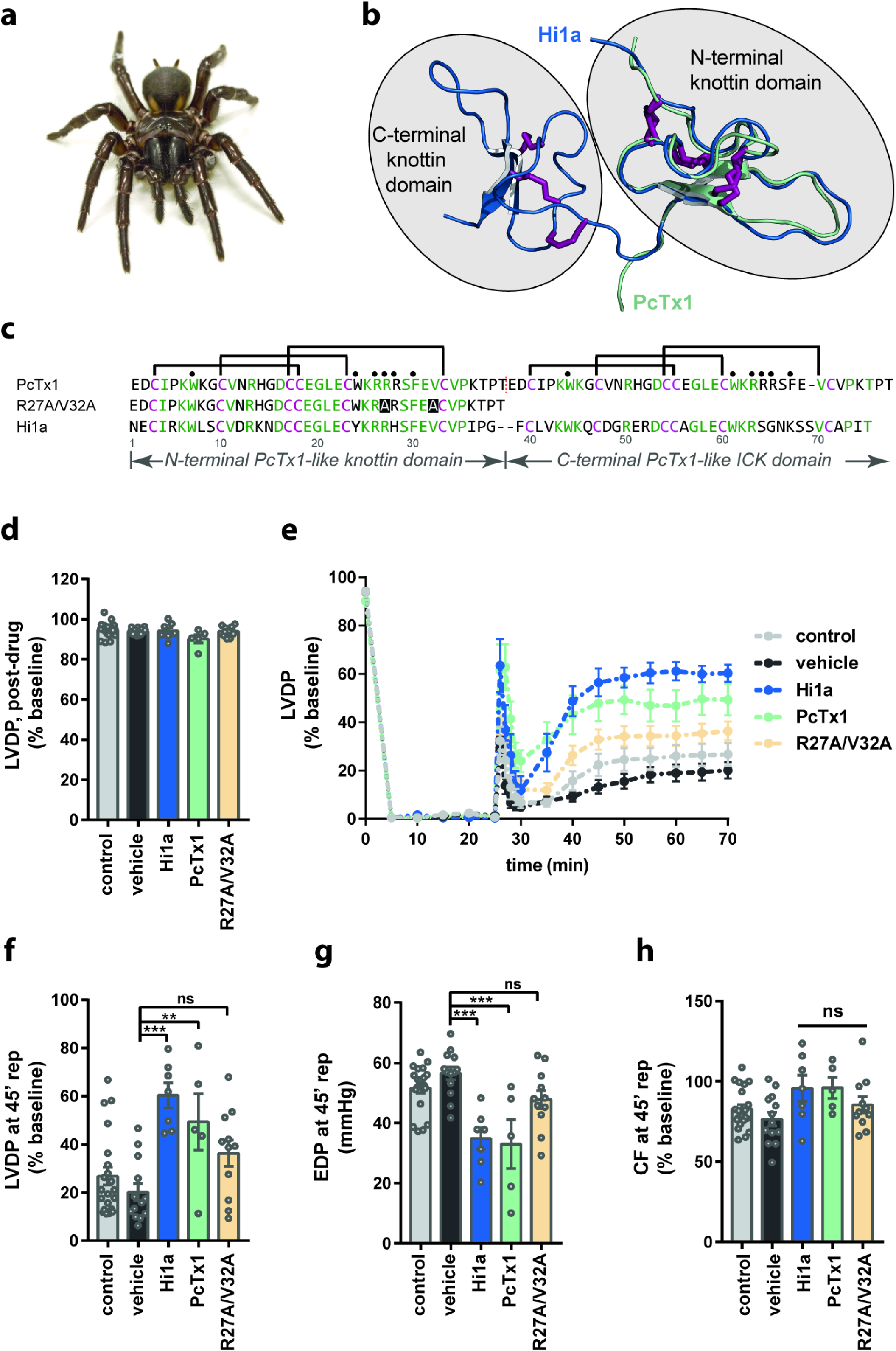
ASIC1a inhibitors protect isolated mouse hearts from IRI. (**a**) Australian funnel-web spider, *Hadronyche infensa*, from which the ASIC1a inhibitor Hi1a was isolated. (**b**) Schematic of the 3D structure of Hi1a (PDB 2N8F^32^) highlighting the two knottin domains. The 3D structure of PcTx1 (green; PDB 2KNI^65^) is overlaid on the N-terminal knottin domain of Hi1a (blue). The disulfide bonds in each structure are shown as maroon tubes. (**c**) Sequence alignment of Hi1a, PcTx1, and the PcTx1-R27A/V32A analogue. Conserved residues are shown in green, except cysteine residues which are highlighted in maroon. Black circles indicate pharmacophore residues of PcTx1. (**d-h**) Langendorff-perfused hearts from adult (12–14 weeks old) male C57BL/6 mice were subjected to 25 min of global ischemia followed by 45 min of reperfusion. Control hearts (no treatment, *n* = 21) were compared to hearts treated with 10 nM Hi1a (*n* = 7), 10 nM PcTx1 (*n* = 5), 10 nM PcTx1-R27A/V32A, *n* = 11), or 0.1% BSA in water (vehicle control, *n* = 13). For treated hearts, vehicle or peptide solution was infused for 10 min prior to ischemia and during the first 15 min of reperfusion. (**d**) Pre-ischemia LVDP, expressed as % baseline, after exposure to vehicle or peptide for 10 min. (**e**) LVDP over time. (**f-h**) Functional parameters measured or calculated after 45 min of reperfusion including (**f**) LVDP (**g**) EDP and (**h**) CF. For LVDP and CF, baseline values were obtained prior to peptide/vehicle infusion, or 10 min prior to onset of ischemia (no-infusion controls). All data are expressed as mean ± SEM. Statistical significance was determined using one-way ANOVA (**p < 0.01, ***p < 0.001).

To examine if these ASIC1a inhibitors are cardioprotective, we assessed tolerance to IRI in Langendorff-perfused isolated mouse hearts with and without peptide treatment. Consistent with genetic ablation of ASIC1a, Hi1a, PcTx1, and PcTx1-R27A/V32A had no effect on baseline contractile function during the first 10 min of peptide infusion prior to ischemia (**Fig. 2d**). Functional tolerance to IRI was greater in hearts exposed to Hi1a or PcTx1 (10 nM) compared to control hearts (**Fig. 2e**). At the end of reperfusion, Hi1a- and PcTx1-treated hearts, but not hearts treated with the PcTx1-R27A/V32A analogue, had markedly improved recovery of LVDP (Hi1a: 60 ± 4%, PcTx1: 49 ± 12%, PcTx1-R27A/V32A: 36 ± 6%) compared to vehicle controls (20 ± 4%) (**Fig. 2f**). Similarly, treatment with Hi1a and PcTx1, but not PcTx1-R27A/V32A, led to reduced EDP after 45 min reperfusion (Hi1a: 35 ± 4 mmHg; PcTx1: 33 ± 8 mmHg; PcTx1-R27A/V32A: 48 ± 3 mmHg) compared to vehicle controls (57 ± 2 mmHg) (**Fig. 2g**). No differences in final CF were observed between groups (**Fig 2h**), although hearts treated with PcTx1, but not Hi1a, displayed significant reactive hyperaemia during early reperfusion, as evidenced by increased CF during the first 5 min of reperfusion (**Supplementary Fig. 3a,b**). Taken together, our data show that ASIC1a inhibitors protect the heart from myocardial IRI and recapitulate the functional benefits of genetic ablation of the channel.

### Preconditioning with Hi1a improves functional recovery of isolated rodent hearts after prolonged hypothermic ischemia

In order to evaluate the cardioprotective effect of ASIC1a inhibition in a model of donor heart preservation, we supplemented standard clinical heart preservation solution (Celsior^38^) with Hi1a during 8 h cold storage of isolated rat hearts. The hearts were stabilised for 10 min by retrograde (Langendorff) perfusion and then switched to working-mode in order to assess cardiac function in the presence of physiological afterload. Following assessment of baseline hemodynamic parameters, hearts were arrested by infusion of cold (2–3°C) Celsior solution with and without 10 nM Hi1a into the coronary circulation. After cardioplegic arrest, hearts were stored under hypothermic conditions for 8 h, followed by reperfusion and functional analysis (**Fig. 3a**). Hearts preserved with Hi1a supplementation had markedly improved recovery of aortic flow (AF) (50 ± 11%) and cardiac output (CO) (57 ± 8%) compared to Celsior-only controls (AF: 8 ± 5%; CO: 24 ± 5%) (**Fig. 3b,c**). These data demonstrate that Hi1a is highly cardioprotective against IRI in a model of donor heart preservation with prolonged cold storage.

**Figure 3.**
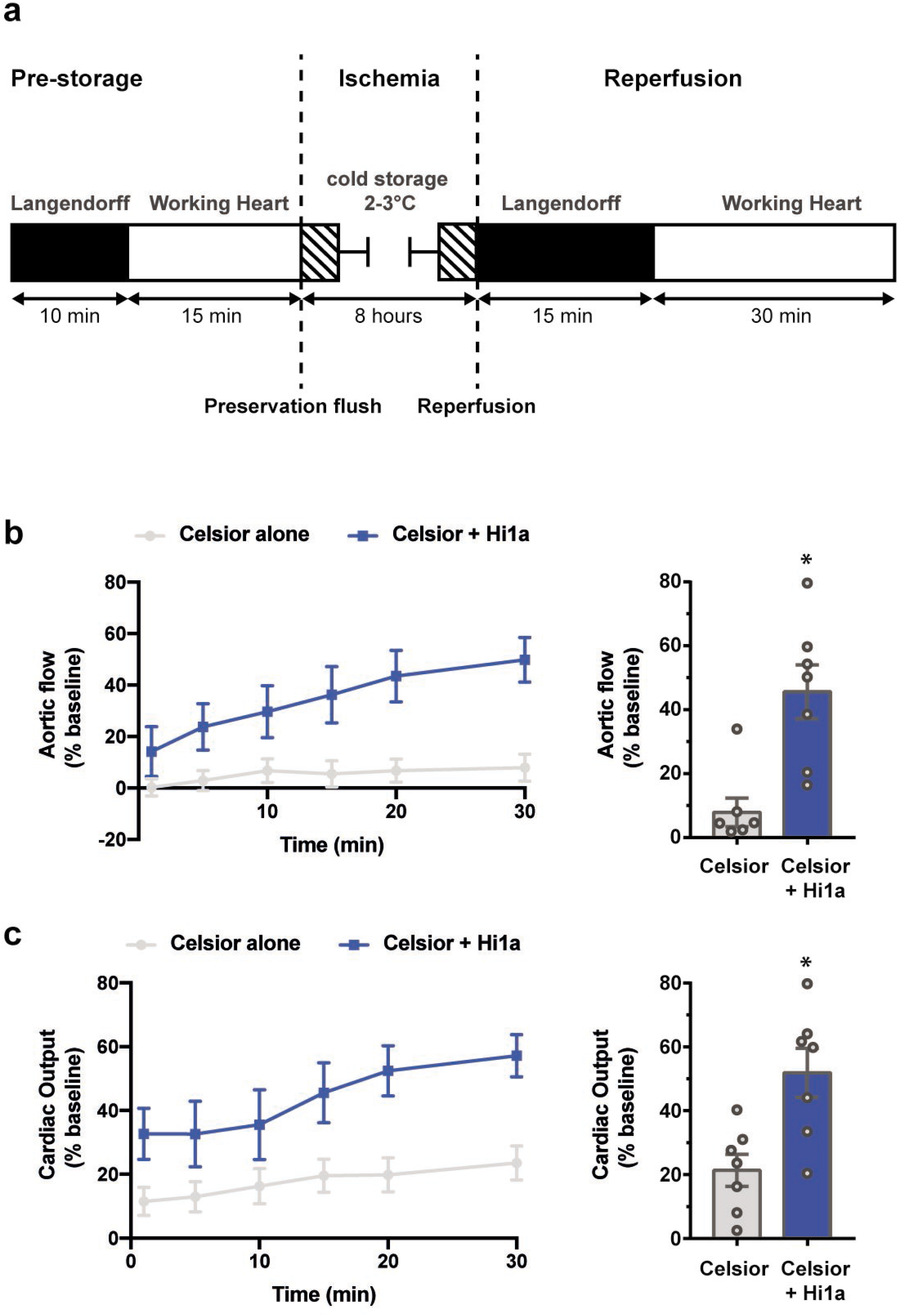
Hi1a protects isolated rat hearts during long-term hypothermic storage. (**a**) Experimental schematic of 8 h cold storage protocol. (**b-c**) Functional parameters assessed throughout 30 min working mode (post-storage reperfusion) and normalized to pre-storage function. (**b**) Aortic flow (% baseline) over time (left) and after 30 min (right). (**c**) Cardiac output (% baseline) over time (left) and after 30 min (right). All data are expressed as mean ± SEM (*n* = 7 per group). Statistical significance was determined using two-tailed unpaired student’s t-test (*p < 0.05).

### Hi1a supplementation improves recovery in a postconditioning model of donation after circulatory death

To determine whether pharmacological inhibition of ASIC1a could recover hearts when delivered subsequent to a profound ischemic injury, we assessed the effect of Hi1a in a rodent model of donation after circulatory death (DCD). Historically, DCD hearts were considered unusable for transplantation due to the severe ischemic injury sustained during organ procurement^39^, but recent studies have shown that successful transplantation of DCD hearts is possible with effective post-conditioning strategies, namely normothermic *ex vivo* perfusion and pharmacological post-conditioning^40–42^. To mimic a clinical DCD withdrawal in rodents, asphyxiation was induced via tracheal ligation, then the hearts were retrieved after a 10-min warm ischemic time (WIT) followed by 2-min stand-off period (after cessation of circulation). The hearts were then flushed with Celsior with or without supplementation, followed by *ex vivo* reperfusion in Langendorff mode for 30 min and then working mode for an additional 30 min (**Fig. 4a**).

**Figure 4.**
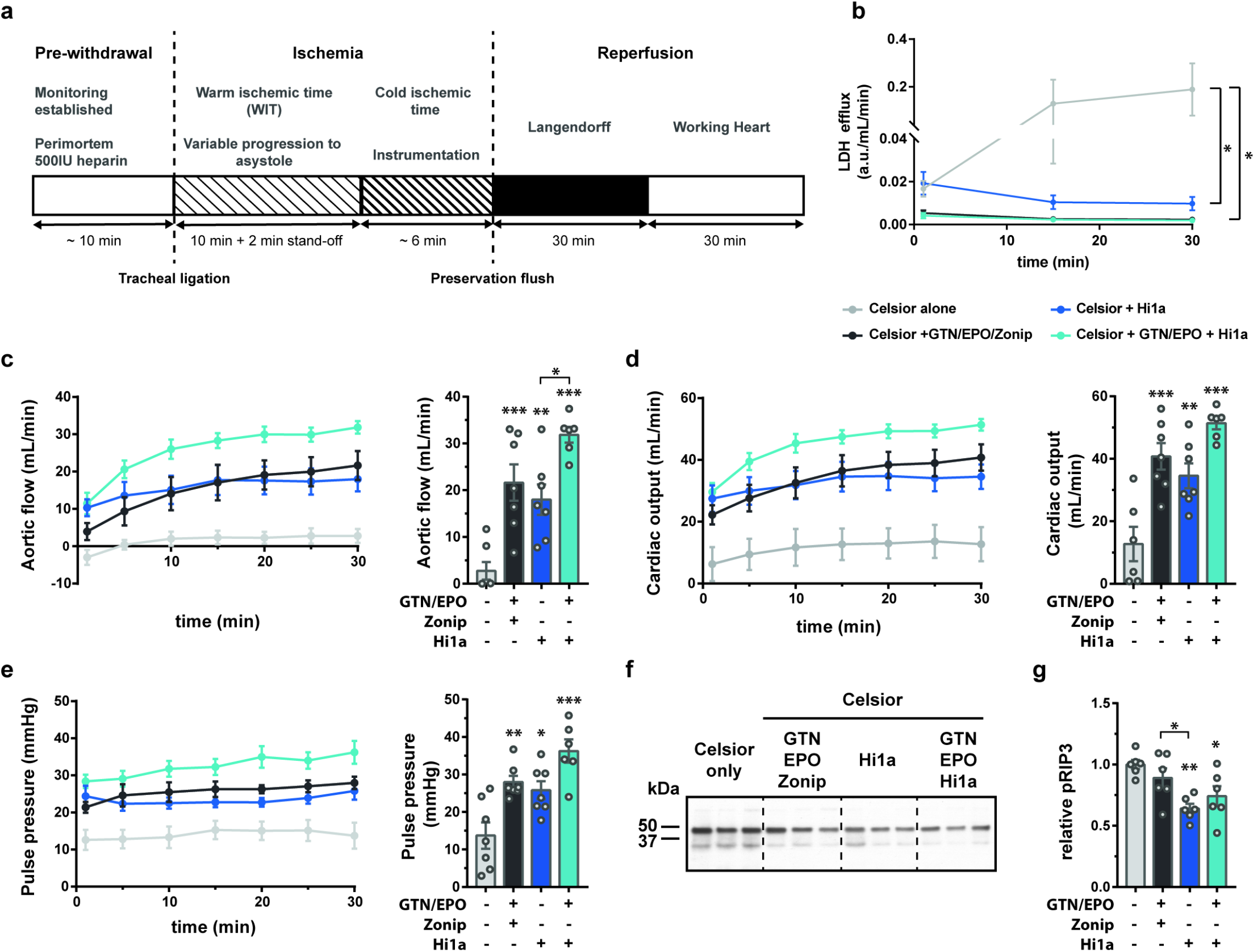
Hi1a protects rat hearts in a preclinical model of DCD. (**a**) Experimental design to mimic clinical DCD withdrawal. (**b-f**) Biochemical and functional characterisation of DCD hearts flushed with preservation solution (Celsior ± supplementation). (**b**) Arbitrary LDH units in the coronary effluent throughout 30 min working mode (normalized to coronary flow rate, LDH a.u./mL/min). (**c-e**) Assessment of heart function with (**c**) AF (mL/min), (**d**) CO (mL/min), and (**e**) PP (mmHg) plotted over time (left) and after 30 min in working mode (right). (**f**) Representative pRIP3 western blot on pRIP3-IP lysates collected at the end of reperfusion. Molecular weight markings (on the left) as determined by BIO-RAD Precision Plus Protein ladder. Individual lanes represent different hearts (*n* = 6 hearts/group with 3 hearts/group/membrane). (**g**) Quantification of pRIP3 relative signal intensity (upper band, ~46 kDa), normalized internally to Celsior-only control. All data are expressed as mean ± SEM (*n* = 6 – 7/group). For all experimental groups, *n* = 6 hearts. Statistical significance was determined using one-way ANOVA (*p < 0.05; **p < 0.01, ***p < 0.001).

Hi1a (10 nM) was tested as a single supplement and in combination with two clinically used supplements, glyceryl trinitrate (GTN) and erythropoietin (EPO)^42^. As a positive control, we also evaluated zoniporide (Zon) combined with GTN and EPO, which we have previously shown activates ischemic postconditioning pathways^41^. To assess the viability of hearts post-DCD withdrawal, functional parameters and cell death (LDH release) were measured throughout working-mode reperfusion. In Celsior-only controls, coronary effluent levels of LDH increased throughout the 30 min assessment period. Allografts supplemented with Hi1a, Hi1a + GTN/EPO, or GTN/EPO/Zon, however, had significantly reduced levels of LDH after 30 min, with LDH release in both triple-supplemented groups comparable to a heart at baseline, prior to ischemic insult (**Fig. 4b**). Functional parameters of recovery including AF, CO, and pulse pressure (PP) were all significantly improved in supplemented groups compared to Celsior alone (**Fig. 4c-e**). AF, in particular, was significantly increased in hearts supplemented with Hi1a combined with GTN and EPO (32 ± 2 mL/min) compared to Hi1a as a single supplement (18 ± 3 mL/min), suggesting an additive effect of Hi1a with current clinically used cardioprotective agents (**Fig. 4c**).

To investigate the therapeutic benefit of Hi1a in the context of preventing cell death in DCD hearts, we performed western blots to examine the levels of phosphorylated RIP3 (pRIP3) in tissue lysates collected at the end of working-mode reperfusion. RIP3 is one of the key regulators of the necroptosis pathway that has been implicated in the response to IRI^43,44^, and specifically to ASIC1a-mediated cell death during cerebral ischemia^45^. Treatment with Hi1a as a single or triple supplement led to reduced levels of pRIP3 following DCD withdrawal and reperfusion (**Fig. 4f**), suggesting that Hi1a blockade of ASIC1a might inhibit necroptosis. Together with our finding that Hi1a improved donor organ viability after prolonged cold storage, the therapeutic efficacy of Hi1a in the DCD model demonstrates significant translational potential for ASIC1a-targeted therapies during clinical donor heart preservation.

### ASIC1a is expressed in the human heart and polymorphisms in its genetic locus are associated with ischemic disease

Analysis of ASIC expression patterns from mRNA-seq and ribo-seq data collected from human left ventricular cardiac tissue^46^ revealed the highest expression of *ASIC1* and *ASIC3* in the human left ventricle and similar levels of *ASIC1* expression in samples from HF and non-HF patients (**Supplementary Fig. 4a,b**).

We next assessed whether natural variation in the genetic locus encoding ASIC1a *(ACCN1)* is associated with ischemic diseases using human population statistical genetics. While fewer than 10% of new molecular entities pass through clinical trials and are approved for therapeutic use, evidence of human genetic association between the gene target and traits sufficiently similar to the indication being treated increase the likelihood of approval to as high as 50%^47^. We utilized summary data from genome-wide association studies (GWAS)^48^ and calculated the statistical significance (using fastBAT^49^) of genetic variants encompassing the *ACCN1* locus with human cardiac and cerebral ischemic phenotypes. These data revealed a significant association between genetic variation in *ACCN1* and major coronary heart disease and MI (P < 0.05) (**Table 2**). We also found that genetic variation in *ACCN1* associates with small-vessel ischemic stroke (p = 3.94 x 10^-3^), but not other stroke subtypes (**Table 2**). Taken together, these data provide evidence that *ACCN1* polymorphisms within the human population are associated with ischemic diseases of the heart and brain.

**Table 2.** Association of polymorphisms in *ACCN1* with ischemic conditions. *ACCN1* gene position: chromosome 12, start: 50451419, end: 50477405)

| Genotype | No. of SNPs | SNP start | SNP end | $\chi^2$ statistic | Mean p-value |
| --- | --- | --- | --- | --- | --- |
| Acute myocardial infarction [1] | 75 | rs11377593 | rs2242507 | 152.445 | 1.36E-02 |
| Major coronary heart disease excluding revascularizations [1] | 75 | rs11377593 | rs2242507 | 199.114 | 1.79E-03 |
| Myocardial infarction [1] | 75 | rs11377593 | rs2242507 | 186.588 | 3.07E-03 |
| Any stroke [2] | 61 | rs78972052 | rs2242507 | 81.8082 | 1.45E-01 |
| Any ischemic stroke [2] | 62 | rs78972052 | rs2242507 | 77.2837 | 1.98E-01 |
| Large artery stroke [2] | 63 | rs78972052 | rs2242507 | 67.2951 | 3.48E-01 |
| Cardioembolic stroke [2] | 61 | rs78972052 | rs2242507 | 71.6309 | 2.52E-01 |
| Small vessel stroke [2] | 64 | rs78972052 | rs2242507 | 147.484 | 3.94E-03 |

The number of SNPs inside the *ACCN1* locus was used to calculate the $\chi^2$ statistic and p-value. Source data: GWAS results from [1] UK Biobank from Neale Lab<sup>48</sup> or [2] Malik et al.<sup>50</sup>.

### ASIC1a inhibition prevents cell death in *in vitro* human models of IRI

To examine whether pharmacological blockade of ASIC1a might provide therapeutic benefit in the context of human tissue, we used human induced pluripotent stem cell-derived cardiomyocytes (hiPSC-CMs). Contractile cardiomyocytes were generated from stem cells using a standard monolayer-based differentiation protocol involving bi-phasic Wnt regulation (**Fig. 5a** and **Supplementary Fig. 5a**). Analysis of mRNA transcripts from single-cell RNA sequencing data over a time course of cardiac differentiation from pluripotency^51^ revealed that *ASIC1, ASIC3*, and *ASIC4* expression increased starting in day-5 cardiac progenitor populations and peaked in day-15 definitive cardiomyocytes. *ASIC1* was the most highly expressed ASIC subtype, with expression levels comparable to other cardiac ion channels such as TRPV1 and NaV1.5 (**Fig. 5b,c**).

**Figure 5.**
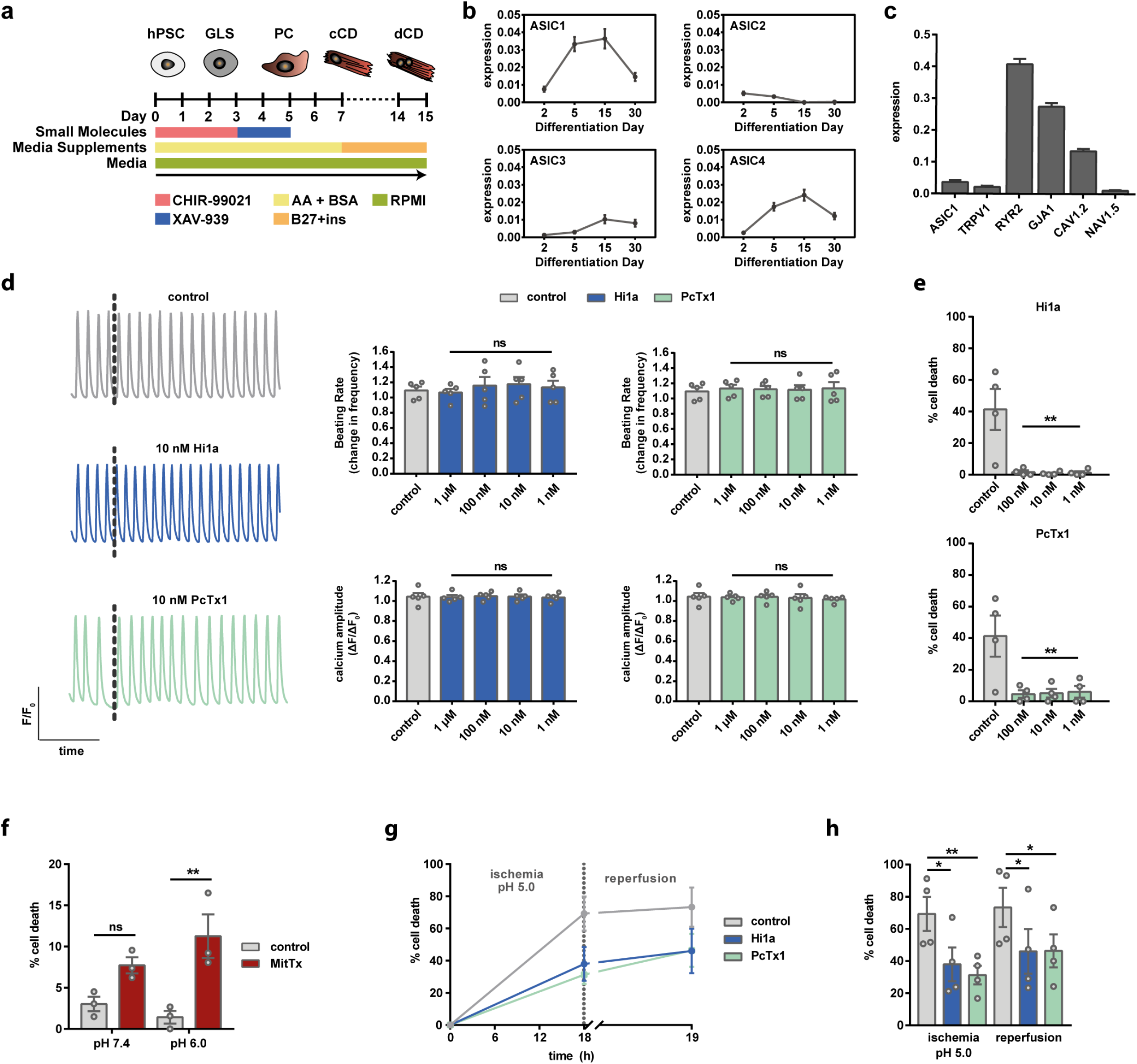
Hi1a protects hiPSC-CMs from ischemic injury. (**a**) Schematic depicting directed differentiation of hiPSC-CMs (adapted from Ref.^51^). (**b-c**) Analysis of published transcriptomic data (scRNAseq) of cardiac differentiation^51^. (**b**) Expression of ASIC1, ASIC2, ASIC3, and ASIC4 at day 0, 2, 5, 15, and 30 of differentiation. (**c**) Gene expression in day-15 hiPSC-CMs. Abbreviations: RYR2 = ryanodine receptor 2; GJA1 = gap junction alpha1 protein (connexin 43). (**d**) Fluorescent imaging of calcium transients (normalized arbitrary fluorescent units (F/F_0_)) before and after Hi1a or PcTx1 addition using a FLIPR Tetra system. Representative traces (black dotted line indicates time of peptide addition) and quantification of spontaneous beat rate and calcium amplitude are shown. Amplitude and beat rate are both expressed as a response over baseline (post-addition measurement normalized to pre-addition measurement). (**e**) Cell death (LDH secretion) analysis after overnight treatment in HBSS pH 5.0 with or without Hi1a (top) or PcTx1 (bottom). (**f**) Cell death (LDH) after overnight treatment with 20 nM MitTx in HBSS pH 7.4 or HBSS pH 6.0. (**g-h**) Cell death (LDH) after *in vitro* IRI with overnight hypoxic (0.5% O_2_) incubation in HBSS pH 5.0 followed by 1 h reperfusion with HBSS pH 7.4 in normoxic conditions. For panels e–h, data are expressed as percent cell death calculated from LDH levels in low (RPMI + B27) and high (1% Triton X-100 in RPMI + B27) controls. All data are expressed as mean ± SEM (*n* = 3–5 biological replicates with 3–6 technical replicates each). Statistical significance was determined using one-way ANOVA (*p < 0.05; **p < 0.01).

Since ASIC1a mediates influx of calcium in addition to sodium^34^, we performed calcium imaging after acute addition of the ASIC1a inhibitors Hi1a and PcTx1 to assess whether ASIC1a inhibition alters physiological electromechanical coupling in cardiomyocytes. Replated hiPSC-CMs were loaded with a fluorescent calcium dye and calcium transients were recorded before and after peptide addition. Neither Hi1a nor PcTx1 altered calcium amplitudes or spontaneous beating rate at any evaluated concentration (**Fig. 5d**).

To assess whether ASIC1a inhibition is cardioprotective in hiPSC-CMs, we evaluated cell death in response to *in vitro* acidosis, induced by culturing hiPSC-CMs in Hank’s buffered sodium salt (HBSS) with pH adjusted to pH 7.4, 6.0, or 5.0. *ASIC1* mRNA expression was not significantly altered by low pH treatment, but significant cell death (> 40%), as assessed by LDH secretion, was observed in populations treated overnight at pH 5.0, with minimal cell death occurring at pH 7.4 or pH 6.0 (**Supplementary Fig. 5b,c**). Treatment with either Hi1a or PcTx1 resulted in nearly complete cardioprotection, even at concentrations as low as 1 nM (**Fig. 5e** and **Supplementary Fig. 5d,e**). To further confirm that ASIC1a plays a direct role in mediating cell death, we treated hiPSC-CMs overnight with 20 nM MitTx, a potent agonist of ASIC1a isolated from snake venom^52^. Consistent with ASIC1a mediating the injury response to cardiac ischemia, treatment of hiPSC-CMs with MitTx resulted in increased cell death at both pH 7.4 and pH 6.0 (**Fig. 5f**). We next evaluated the cardioprotective effect of Hi1a and PcTx1 in an *in vitro* model of ischemia/acidosis with reperfusion. To mimic ischemia/acidosis *in vitro*, hiPSC-CMs were incubated overnight in combined hypoxic (0.5% O_2_) and acidic (pH 5.0) conditions with or without peptide. After 18 h of incubation, the low-pH medium was replaced with medium at physiological pH 7.4 with or without peptide, and the cells were incubated for an additional hour under normoxic conditions. Significant cell death was observed in control hiPSC-CMs, but this was blocked by either 10 nM Hi1a or PcTx1 (**Fig. 5g,h**). These data suggest that ASIC1a mediates cell death responses in human cardiomyocytes and that pharmacological inhibition of ASIC1a confers significant protection against ischemia-induced cell stress.

## Discussion

Despite decades of research and promising preclinical data, no pharmacological strategies have led to robust improvement in clinical outcomes for ischemic injuries of the heart^10^. In this study, we showed for the first time that ASIC1a exacerbates cardiomyocyte injury response to ischemia. Our findings are consistent with prior research demonstrating the importance of ASIC1a during ischemic stroke^30–32^, and they extend the potential therapeutic applications of ASIC1a-targeted therapies to myocardial IRI.

Although ASICs, and in particular ASIC1a, are generally associated with the central and peripheral nervous systems^53^, our expression analyses revealed that ASIC1 is present in both rodent and human cardiomyocytes, and is not differentially regulated at the transcriptional level as a result of cellular ischemia *in vitro* or *in vivo*^33^. Our evidence of ASIC1a expression in the heart is supported by recent studies that identified ASIC-like currents in hiPSC-CMs^54^ and rat cardiomyocytes^23^. In neurons, there are several possible mechanisms by which ASIC1a activation leads to cell death including calcium overload^55^, regulation of mitochondrial permeability transition (MPT) pores^56^, and the initiation of necroptosis through direct interaction with, and activation of, RIPK1^45^. ASIC1a-induced cell death has not previously been described in cardiomyocytes, but there is evidence that ASIC1a, along with TRPV1, mediates calcium influx in response to extracellular acidosis^23^. Importantly, given the potential for ASIC1a to transport calcium, we showed that genetic ablation or pharmacological inhibition of ASIC1a does not alter normal electromechanical coupling in the heart *in vitro* or *in vivo*.

Given that drug targets with genetic evidence of disease association are twice as likely to succeed in clinical trials and lead to approved drug candidates^47^, our GWAS analysis of the association between genetic variation in the *ACCN1* locus and ischemic disease points towards the translational potential of ASIC1a-targeted therapies. Such therapies have the potential to impact a broad scope of clinical applications since myocardial IRI is evident in many cardiovascular complications such as acute MI, out-of-hospital cardiac arrest (OHCA), cardiopulmonary bypass, and heart preservation for transplantation^7^. Despite intensive research into therapies aimed at mitigating myocardial IRI in the context of acute MI, none have translated into clinical practice. Drugs that prevent calcium overload by targeting NHE showed translational promise in preclinical studies^57^, but they ultimately failed in Phase III clinical trials (The EXPEDITION Trial) due to cerebrovascular complications^19^. Nonetheless, this trial provided strong clinical evidence that myocardial IRI is a threat during coronary artery bypass graft surgery and that NHE inhibitors can reduce post-operative MI^19^.

In general, there is a strong association and co-morbidity between stroke and cardiac arrest, with 88% of patients suffering some form of cardiovascular complication within 4 weeks of an ischemic stroke^58^. Likewise, cardiac interventions such as transcatheter aortic valve implantation (TAVI) can cause cerebrovascular complications^59^. We previously used a rat model of ischemic stroke to show that a single dose of Hi1a (2 ng/kg i.c.v) massively reduced infarct size and improved neurological and motor performance even when administered up to 8 h after stroke onset^32^. No other drugs have been shown to provide significant protection beyond 4 h after stroke onset^60^. Thus, ASIC1a inhibitors might provide a potential major advantage for clinical use due to their ability to protect both the heart and the brain from IRI, which is particularly relevant for OHCA, for which there is both cardiac and cerebral manifestations.

ASIC1a-targeted therapies appear to have significant potential in heart transplantation applications. The predominant source of donor hearts is brain dead (BD) donors, but the recent introduction of DCD hearts has increased the donor pool^42^. The viability of usable BD and DCD hearts, however, continues to be adversely affected by unavoidable injury to the heart that occurs during organ procurement and storage, as well as reperfusion injury in the recipient. Indeed, the relative risk of one-year morbidity is strongly correlated with organ ischemic time^7,61^. In general, primary graft dysfunction is a major risk for patients in the first 30 days after transplantation and accounts for 66% of patient deaths, of which 48% are due to IRI^62^. Thus, there is an unmet need for effective, early interventions that reduce injury to the graft and reduce the risk of primary graft dysfunction in the recipient. Our direct comparison between Hi1a and supplements in current clinical use provide strong evidence for the clinical utility of Hi1a for increasing the pool of viable donor hearts for transplantation.

Venom-derived molecules have proven to be a source of novel therapeutics, particularly in the cardiovascular field. Examples include the antihypertensive drug captopril^63^ derived from the venom of a Brazilian viper, and eptifibatide and tirofiban, antiplatelet drugs that are used to treat acute coronary syndromes^64^. Disulfide-rich knottin peptides such as Hi1a are typically highly stable in biological fluids, and ziconitide, a knottin peptide derived from cone snail venom, is an FDA-approved analgesic^64^. For this reason, Hi1a is ideally suited for clinical use as an adjuvant to standard-of-care heart preservation solutions for heart transplant or in combination with mechanical procedures such as percutaneous coronary intervention for myocardial infarction.

In summary, we have shown that ASIC1a is a critical mediator of cell death during cardiac ischemia and that pharmacological inhibition of ASIC1a affords potent cardioprotection across rodent and human models of cardiac IRI. These findings suggest that ASIC1a inhibitors have the potential to significantly impact the public health burden of cardiovascular disease, and result in improved outcomes for patients with cardiac ischemic injuries.

## Acknowledgements

We gratefully acknowledge funding support from The University of Queensland (UQ) Strategic Funding Research Initiatives (to NJP, GFK, JFF, JEH, ERP, MER, WGT), the Australian National Heart Foundation (Grant 101889 to NJP), the Whitaker International Program (Scholar Grant to MAR), the American Australian Association (Grant 441 to MAR), the St Vincent’s Clinic Foundation (to PSM), the Australian National Health & Medical Research Council (Program Grant 1074386 to PSM), the Australian Research Council (Discovery Early Career Researcher Award DE180100976 to GCP), and the Australian National Health & Medical Research Council (Principal Research Fellowship APP1136889 and Development Grant APP1158521 to GFK, Project Grant APP1085996 to WGT). Thanks to the Integrated Physiology Facility and Melanie Flint at UQ for technical support and assistance. Microscopy was performed at the Australian Cancer Research Foundation (ACRF)/Institute for Molecular Bioscience Cancer Biology Imaging Facility, which was established with the support from ACRF. We thank Andrew Harvey for assistance in assessing the landscape of pharmaceutical development of ASIC1a inhibitors.

## Author Contributions

**M.A.R.** designed, performed, and analysed the experiments presented in **Figs 1**, **2**, and **5**, **Table 1**, and **Supplementary Figs 1**, **3**, and **5**, and wrote the manuscript. **S.E.S.** designed, performed, and analysed the experiments described in **Figs 3** and **4** and wrote the manuscript. **N.J.S.** produced, purified, and performed QC analyses for peptides used in **Figs 2 – 5** and **Supplementary Figs 3** and **5**. **L.G.**, **M.H.**, and **J.E.V.** helped design, perform, and interpret experiments described in **Figs 3** and **4**. **M.E.R.**, **W.G.T.**, **J.N.P.**, **J.Y.S**., and **L.E.S.H.** conceived ideas and provided insights towards interpretation of results shown in **Figs 1** and **2**. **H.S.C.** and **X.C.** helped perform the experiments described in **Fig. 5** and **Supplementary Fig. 5**. **Y.S**. performed computational analysis presented in **Supplementary Fig. 4**. **J.E.H, G.A.Q,** and **E.R.P.** generated data and analysed ASIC isoform expression in the heart in **Fig 1a. G.C.P** performed GWAS analysis and interpretation presented in **Table 2**. **J.F.F** helped conceive the heart transplant studies and provided intellectual and financial support. **B.R., R.J.H, S.M.,** and **S.P** generated and validated the ASIC1a knockout mouse strain used for studies presented in **Fig. 1** and **Supplementary Fig. 2**. **G.F.K.** conceived of experimental objectives, provided intellectual and financial support, and directly supervised work related to peptide production utilized across all aspects of the study. **P.S.M** conceived of experimental objectives, provided intellectual and financial support, and directly supervised work related to heart transplant modelling. **N.J.P.** conceived of experimental objectives, provided intellectual and financial support, directly supervised work related to mouse IRI, GWAS analysis, human pluripotent stem cell modelling, and oversaw programmatic decisions regarding study design and completion in close collaboration with **P.S.M** and **G.F.K**. All authors contributed to revisions and final preparation of the manuscript.

## Online Methods

### Animals

All animals used in this study received humane care in compliance with National Health and Medical Research Council (Australia) guidelines and the *Guide for the Care and Use of Laboratory Animals* (National Institute of Health, Bethesda, MD). All experiments were performed in accordance with protocols approved by the Animal Ethics Committee of The University of Queensland (IMB/171/18) and Garvan Institute of Medical Research (16/38). To test the therapeutic efficacy of ASIC1a inhibitors, male C57BL/6 mice (Langendorff IRI experiments) or male Wistar rats (donor organ preservation experiments) were purchased from the Animal Resource Centre (Canning Vale, Western Australia).

For genetic ablation studies, ASIC1a^-/-^ mice were generated at the Australian Phenomics Facility. CRISPR/Cas9 technology was used to specifically target the mouse ASIC1a sequence. Use of guide RNAs (gRNA) with following sequences CCGAGGAGGAGGAGGTGGGTGGT and GTACCATGCTGGGGAACTGCTGG, resulted in single nucleotide deletions within both targeted regions, at positions 22 and 341 (NM_009597.2(ASIC1_v001):c.22del and NM_009597.2(ASIC1_v001):c.341del). These deletions predicted a disrupted ASIC1a protein sequence (p.Glu8Argfs*9 and p.Leu114Argfs*94, respectively). The founder mouse was backcrossed to C57BL/6 background. Both hetero- and homozygous mice were viable and showed no obvious phenotype. Total RNA was isolated from brain tissue using Trizol Reagent (ThermoFisher Scientific, Massachusetts, USA), and contaminant genomic DNA was removed with DNA-free reagents (Ambion/Life Technologies, Austin, USA). Primer sequences designed to distinguish between ASIC1a and ASIC1b transcripts were used to determine gene expression levels in ASIC1a KO and WT animals^66^. Primers sequences used in this study were ASIC1a forward 5’-CTGTACCATGCTGGGGAACT-3’ and reverse 5’-GCTGCTTTTCATCAGCCATC-3’; Asiclb forward 5’-TGCCAGCCATGTCTTTGTG-3’ and reverse 5’-CACAGGAAGGCACCCAGT-3’ and RPL32 (for sample normalization) forward 5’-GAGGTGCTGCTGATGTGC-3’ and reverse 5’-GGCGTTGGGATTGGTGACT-3’. For quantitative real-time (qRT)-PCR, oligo-dT primed cDNA was synthesized from 500 ng of total RNA using Murine Moloney Leukaemia Virus reverse transcriptase (Promega, USA). qRT-PCR was performed using a ViiA Real-Time PCR System (Applied Biosystems, California, USA) using SYBR green master mix (Promega, USA) according to manufacturer protocols. Relative ASIC1a and ASIC1b gene expression values were obtained from ASIC1a^-/-^ mice and WT (ASIC1a^+/+^) mice (calibrator) by normalization to the reference gene RPL32 using the 2-ΔΔCt method, where 2-ΔΔCt = ΔCt sample – ΔCt calibrator.

### IRI in Langendorff-perfused mouse hearts

Isolated hearts were assessed for tolerance to IRI as previously described^67,68^. Mice (12–14 weeks) were anesthetized via an intraperitoneal injection of 10 mg/mL ketamine and 1.6 mg/mL xylazil. The heart was excised via thoracotomy and the aorta cannulated. The hearts were retrogradely perfused under constant hydrostatic pressure (80 mmHg) with oxygenated (95% O_2_; 5% CO_2_) modified Krebs-Henseleit buffer (composition in mM: 119 NaCl, 22 NaHCO_3_, 4.7 KCl, 1.2 MgCl_2_, 1.2 KH_2_PO_4_, 0.5 EDTA, 1.85 CaCl_2_, 11 d-(+)-glucose, 2 Na^+^ pyruvate, all from Sigma-Aldrich). Temperature was continuously measured with thermocouple needle microprobes (SDR Scientific) and maintained at 37°C with circulating water baths. Contractile function was measured via a fluid-filled balloon inserted in the left ventricle and connected to a pressure transducer (ADInstruments Pty Ltd.). Coronary flow was measured with an in-line Doppler flow probe positioned above the aortic cannulae (Transonic Systems Inc.). All functional data were recorded on a four-channel MacLab system (ADInstruments Pty Ltd.). Following 15–30 min of equilibration, hearts were switched to ventricular pacing at 420 beats/min (BPM) using a SD9 stimulator (Grass Instruments, Inc.). Baseline measurements were made for 10–20 min followed by 25 min of global normothermic ischemia and 45 min of reperfusion. Pacing was stopped during ischemia and resumed after 15 min of reperfusion.

For peptide-treated experimental groups, 1 μM peptide solution was infused with a syringe pump (World Precision Instruments, LLC.) into the buffer, directly upstream of the cannula, at a rate of 1% of CF, for a final working concentration of 10 nM. Peptide was infused for 10 min prior to the onset of ischemia and during the first 15 min of reperfusion. To assess cell death, effluent was collected at 2 and 45 min after the start of reperfusion. Effluent levels of LDH were measured using a cytotoxicity detection kit (Roche). Normalized absorbance values (absorbance at 492 nm minus absorbance at 690 nm) were measured using a PHERAStar FS microplate reader (BMG Labtech). Standard curves were generated with bovine LDH (Sigma-Aldrich) and used to convert sample absorbance values to units of LDH. Data were normalized to CF and heart weight and then expressed as U/min/g. For all analyses, hearts were excluded if they met the following criteria: abnormally high coronary flow (> 8 mL/min), delayed onset of ischemic contracture (time to onset of ischemic contracture (TOIC) > 20 min), poor contractile function after equilibration (significant arrhythmias and/or left ventricular systolic pressure < 80 mmHg), or technical issues with the perfusion rig.

### Isolated working model of donor heart preservation after prolonged cold storage

We used an isolated working heart model of donor heart preservation that we previously described^69–71^. Male Wister rats (325–395 g) were anesthetized with an intraperitoneal injection of ketamine (80 mg/kg; Cenvet, Australia) and xylazine (10 mg/kg; Provet, Australia), with supplemental doses given as necessary until adequate anaesthesia was achieved. An incision was made in the abdomen of the animal and a bolus injection of 300 IU of heparin administered into the renal vein. A sternotomy was performed and the heart was excised and arrested by immersion in ice-cold Krebs-Henseleit buffer (composition in mM: 118 NaCl, 4.7 KCl, 1.2 MgSO_4_, 1.2 KH_2_PO_4_, 25 NaHCO_3_, 1.4 CaCl_2_, 11 glucose, all from Sigma-Aldrich). Following aortic cannulation, hearts were stabilised for 10 min by retrograde (Langendorff) perfusion with Krebs-Henseleit buffer (37°C), at a hydrostatic pressure of 100 cm H_2_O. During this period, the left atrial appendage was cannulated. This nonworking Langendorff system was converted to a working preparation by switching the supply of perfusate from the aorta to the left atrial cannula at a hydrostatic pressure of 15 cm H_2_O (preload). The working heart ejected perfusate into the aortic cannula against the maintained fixed hydrostatic pressure of 100 cm H_2_O (afterload). Aortic pressure was monitored via a side arm of the aortic cannula with a pressure transducer (Ohmeda, Pty Ltd., Singapore). Aortic flow (AF) was measured by an in-line flowmeter (Transonics Instruments Inc., Ithaca, NY). Aortic pressure and flow were recorded using PowerLab/4e (ADInstruments Pty Ltd, Sydney, Australia) with heart rate (HR) calculated from the flow trace. Coronary flow (CF) was measured by timed collection of effluent draining from the heart via a small incision in the pulmonary artery (PA). Cardiac output (CO) was calculated from the sum of AF and CF.

**Figure 3a** illustrates the experimental timeline. The heart was maintained in working mode for 15 min, at which time functional parameters (AF, CF, CO and HR) were recorded and defined as the baseline value. Any hearts having a baseline AF < 35 mL/min, HR < 200 bpm, or CF < 10 mL/min were excluded at this stage. The heart was arrested by infusion of a preservation flush containing Celsior solution (2–3°C) ± 10 nM Hi1a via the aortic cannula (60 cm H_2_O infusion pressure over 3 min), immersed in 100 mL Celsior ± 10 nM Hi1a, and kept on ice for 8 h. The heart was remounted on the perfusion apparatus and perfused with fresh Krebs-Henseleit buffer for 15 min in Langendorff mode, followed by 30 min in working mode. Celsior solution was obtained from the Institut Georges Lopez (Lissieu, France). Cardiac functional indices after 30 min of working-mode reperfusion were expressed as percent recovery of baseline values.

### Donor heart preservation following circulatory death in rodents

The experimental model of donation after circulatory death (DCD) in rodents (**Fig. 4a**) was developed by our laboratory to closely mimic the events during withdrawal of life support (WLS) in the clinical scenario. Male Wister rats (325–395 g) were anesthetized with an intraperitoneal injection of ketamine (80 mg/kg; Cenvet, Australia) and xylazine (10 mg/kg; Provet, Australia), with supplemental doses given as necessary until adequate anaesthesia was achieved. Non-invasive monitoring was established, consisting of measurement of pulse oximetry (Rad-5v, Masimo Inc., Irvine, CA) and continuous electrocardiogram (P55 AC pre-amplifier, Grass Instrument Co., RI). A midline incision was made in the neck of the animal to expose the right carotid artery, which was cannulated for invasive monitoring of arterial pressure via a pressure transducer (Ohmeda, Pty Ltd., Singapore). Once baseline monitoring parameters were recorded, a dose of 500 IU for antemortem heparin was delivered via a side port in the arterial pressure monitoring line. The animal was subjected to WLS via asphyxiation. A fixed warm ischemic time of 10 min followed by a 2-min stand-off period (to mimic Australian federal regulations regarding DCD heart retrieval^42^) was chosen. Upon completion of the stand-off period, the heart and lungs were swiftly excised *en bloc* and immersed in ice-cold Krebs-Henseleit buffer. The aorta and left atrium were fashioned for instrumentation on an *ex-vivo* Langendorff perfusion apparatus. Once the aorta was cannulated onto the *ex-vivo* circuit, the pulmonary veins were rapidly ligated, and the preservation flush (containing Celsior solution (2–3°C) ± supplements) was commenced via the aortic cannula at a flow rate of 15–20 mL/min up to a total volume of 100 mL. The following concentrations of supplements were used: Hi1a: 10 nM; GTN: 100 mg/L; EPO: 5 U/mL; zoniporide: 1 μM. Left atriotomy was performed to prevent left ventricle distension and to facilitate cannulation on the *ex-vivo* apparatus for functional assessment during the working phase of the experiment.

Following the preservation flush, Langendorff reperfusion was commenced for a period of 30 min, using oxygenated Krebs-Henseleit buffer (37°C) to allow stabilization and recovery of the heart. As previously described, the apparatus was converted to a working mode via switching the supply of perfusate from the aortic to the left atrial cannula at a hydrostatic pressure of 15 cm H_2_O (preload). Functional parameters (AF, CO, PP and HR) were measured continuously, and CF measured at 5-min intervals, during working mode. Coronary effluent and left ventricular free-wall samples were collected as per the protocol for the isolated working heart model.

To assess cell death throughout working mode, effluent LDH levels were measured as an index of structural cardiomyocyte injury using the TOX 7 assay kit (Sigma-Aldrich). Duplicate 75 μl aliquots were assayed according to the manufacturer’s instructions. The resulting absorbance of the tetrazolium product was measured at 492 nm using a 96-well plate reader and expressed as arbitrary units, normalised to coronary flow per minute.

### Peptide production

Recombinant Hi1a was produced by expression as a His_6_-maltose-binding protein (MBP) fusion protein in the periplasm of *Escherichia coli* as previously described^32^, but with optimisation of the expression and purification conditions to improve the yield of Hi1a. Briefly, *E. coli* BL21(DE3) cells were grown at 30°C in Terrific Broth until the optical density at 600 nm reached 2.0–2.5, at which point the temperature was reduced to 17°C and after a 15 min delay expression of the Hi1a fusion protein was induced with 1 mM IPTG. Cells were harvested after a further 21 h growth. PcTx1 and the PcTx1 - R27A/V32A analogue were produced in the same manner with minor modifications to the expression protocol. *E. coli* were grown at 37°C for the entire expression period and harvested approximately 4–5 h after induction.

Cell pellets were resuspended in 50 mM Tris, 300 mM NaCl, 5% glycerol, 15 mM imidazole (pH 8.3 for Hi1a or pH 8.0 for PcTx1), and the cells were lysed by high-pressure cell disruption (Constant Systems Limited, United Kingdom). The His_6_-MBP-peptide fusion proteins were purified from the clarified soluble lysate over a nickel affinity resin. The resin was washed with the same buffer to elute weakly bound proteins before eluting the Hi1a fusion protein with the same buffer containing 300 mM imidazole. Fusion proteins were exchanged into low (< 30 mM) imidazole buffer using an Amicon Ultra-15 centrifugal concentrators (Merck Millipore, Germany) with a 30 kDa molecular weight cut-off in preparation for liberation of Hi1a from the fusion tag using tobacco etch virus (TEV) protease. The fusion proteins were cleaved in redox buffer containing 3 mM reduced glutathione and 0.3 mM oxidised glutathione at pH 8.3 for Hi1a or pH 8.0 for PcTx1, using ~1 mg of TEV protease per 50 mg of fusion protein. For Hi1a the cleavage reaction was allowed to proceed at 4°C over 3–6 days. For the PcTx1 peptides, cleavage was performed at room temperature for a minimum of 16 h. The recombinant peptides each contain a nonnative N-terminal serine residue, which is a vestige of the TEV protease recognition site. The released peptides were purified from the cleavage reaction solutions using reverse phase high-performance liquid chromatography.

The mass of the purified peptides were confirmed by electrospray-ionisation mass spectrometry and pure peptides were lyophilised prior to confirmation of ASIC1a inhibitory activity using two electrode voltage clamp electrophysiology, as described previously for both Hi1a^32^ and PcTx1^30,65,72^. Unless otherwise noted, lyophilized stocks of peptide were reconstituted in sterile deionised water prior to use.

### Western blot

Left ventricular tissue samples (~60 mg) from hearts (N = 6/group) collected at the end of the DCD protocol were homogenized in ice-cold lysis buffer (150 mM NaCl, 50 mM Tris-HCl, 1% Triton X-100, 1 mM sodium orthovanadate, 1 mM glycerophosphate, 5 mM dithiothreitol (DTT), 15% Roche cocktail protease inhibitors; pH 7.4). Samples were centrifuged at 10,000 rpm (9,400 *g*) for 5 min at 4°C. The protein concentration of each lysate was measured using a Bradford assay kit (Pierce Biotechnology, Rockford, IL), with bovine serum albumin used as standard. If required, immunoprecipitation (IP) was performed (see below). Protein samples (40 μg) were heated at 95°C in Laemmli sample loading buffer for 10 min and electrophoretically separated on 4–20% graduated precast gels (Bio-Rad) and transferred to polyvinylidene difluoride membranes. After blocking with 5% non-fat dry milk in Tris-buffered saline (TBS) containing 0.1% Tween-20, membranes were incubated overnight at 4°C with primary antibody (pRIP3 Ser227, Cell Signalling Technology, USA). Membranes were incubated with secondary anti-rabbit (1:2000 in 5% non-fat dry milk + TBS-T) or anti-mouse (1:5000 in 1% BSA + TBS-T) IgG conjugated to horseradish peroxidase (GE healthcare, Ryadalmere, Australia) for 1 h at 25°C. Protein bands were visualized using SuperSignal West Pico Chemiluminescent Substrate (Life Technologies, Scoresby, VIC, Australia), digitized, and quantified using Image J software (Version 1.52a, National Institute of Health, USA). Within each blot, the signal intensity from each lane was normalized to the Celsior-only group average from the same membrane. Blots were repeated to confirm results. All antibodies, unless otherwise stated, were sourced from Cell Signalling Technology (Beverly, MA).

### Immunoprecipitation

Lysate samples from individual hearts prepared as detailed above were centrifuged for 10 min at 10,000 *g* (4°C). Lysates were diluted with lysis buffer without DTT to make a final volume of 0.5 mL containing 1 mg of total proteins (1 μg/μl). Pre-clearing of the samples was performed by adding 60 μl of 50% Protein A agarose beads to 1 mL of diluted lysate from above. Samples were tumbled for 30 min at 4°C in a tube rotator, prior to being centrifuged for 5 min at 1,000 *g* (4°C), and supernatants were collected for immunoprecipitation. The cleared lysates were divided into two 400-μl aliquots and 1:100 specific primary antibody added to form the Ag/Ab complex (pRIP3 Ser227, Cell Signalling Technology, USA). 50 μl of 50% Protein A beads were added followed by gentle rocking for 2 h at 4°C. Following centrifugation for 30 s at 10,000 *g*, 4°C, the supernatant (containing unbound proteins) was aspirated, and either discarded or snap frozen at −80°C for sequential immunoprecipitation of other antigens. The remaining pellet was washed five times with 500 μl of ice-cold lysis buffer. For each wash, the beads were re-suspended by inversion and centrifuged for 30 s at 1000 *g* at 4°C, prior to the supernatant being aspirated and discarded. The final pellet was resuspended with 50 μl of 2 x SDS sample buffer without β-mercaptoethanol. The samples were mixed vigorously and heat denatured for 5 min at 95°C. Following a 30 s centrifugation at 10,000 *g*, the supernatants were transferred to new vials and β-mercaptoethanol was added. The prepared samples were loaded on 4–20% graduated precast gels (Bio-Rad) and analysed by Western blotting as detailed above.

### Generation of cardiomyocytes from human induced pluripotent stem cells

All hiPSC studies were performed with consent from The University of Queensland’s Institutional Human Research Ethics approval (HREC#: 2015001434). Cardiomyocytes generated in this study were derived from the WTC-11 hiPSC line (Gladstone Institute of Cardiovascular Disease, UCSF)^73,74^. Undifferentiated hiPSCs were maintained on Vitronectin XF (5 μg/mL, Stem Cell Technologies) coated tissue culture dishes as per manufacturer recommendation with either mTeSR medium with supplementation or mTeSR PLUS medium with supplementation (Stem Cell Technologies). Contractile cardiomyocytes were differentiated using a high-density monolayer format as previously described^51^. On day 1 of differentiation, hiPSCs were dissociated with 0.5 mM EDTA solution supplemented with 1.1 mM D-glucose. Single-cell suspensions were plated at a density of 1.2 x 10^5^ cells/cm^2^ and cultured overnight in mTeSR medium supplemented with 10 μM Y-27632 dihydrochloride (Stem Cell Technologies). Once the monolayer reached approximately 80% confluence (usually the following day), differentiation was induced (day 0). The cells were quickly washed with phosphate-buffered saline (PBS) followed by a change in medium to RPMI (ThermoFisher) containing 3 μM CHIR99021 (Stem Cell Technologies), 500 μg/mL BSA (Sigma Aldrich), and 213 μg/mL ascorbic acid (Sigma Aldrich). After 3 days of culture, the medium was exchanged to RPMI containing 500 μg/mL BSA, 213 μg/mL ascorbic acid, and 5 μM Xav-939 (Stem Cell Technologies). On day 5, the medium was replaced with RPMI containing BSA and ascorbic acid as on day 3. Starting on day 7, the cells were fed every other day with RPMI containing 1x B27 supplement with insulin (Life Technologies). Spontaneous beating was typically observed between days 9 and 11 of differentiation.

### *In vitro* ischemia-acidosis injury model with hiPSC-CMs

Differentiated cardiomyocytes were replated on either day 15 or day 17 of differentiation for *in vitro* ischemia/acidosis assays. At the time of replating, a subset of cells (~500,000) was set aside for flow cytometry analysis of cardiomyocyte purity (see below). For all experiments, only cell preparations with greater than 80% sarcomeric α-actinin-positive cardiomyocytes were used. After re-plating, the cells were maintained for an additional 7 days in RPMI + B27. To prepare media for ischemia/acidosis injury, 10x HBSS without sodium bicarbonate (Sigma) was diluted to 1x concentration in sterile tissue culture-grade water. Solutions were buffered with either 12 mM HEPES (for pH 7.4 media, Sigma Aldrich) or 12 mM MES (for pH < 6.5, Sigma Aldrich) and the pH adjusted accordingly with 1 M NaOH. The medium was sterile filtered with 0.22 μm syringe filters (Millipore). Unless otherwise noted, the replated cells were treated overnight (18 h) in HBSS with or without peptide under either normoxic (~18.5% O_2_; 5% CO_2_) or hypoxic (0.5% O_2_; 5% CO_2_) culture conditions. For reperfusion experiments, the medium was replaced with HBSS pH 7.4 (with or without peptide) after overnight incubation and cultured for 1 h under in normoxic conditions.

To assess cell death, the supernatant was collected and LDH levels were measured using a cytotoxicity detection kit (Roche). For all cell culture experiments, percent cell death was calculated using low and high controls. For low control (LC), cardiomyocytes were cultured overnight in standard culture media (RPMI + B27). For high control (HC), cells were cultured in RPMI + B27 containing 1% Triton X-100 (Sigma-Aldrich).

### Flow cytometry

To assess the cardiomyocyte purity of differentiated cell populations, cells were fixed with 4% paraformaldehyde (Sigma Aldrich), permeabilized in 0.75% saponin (Sigma Aldrich), and labelled with Phycoerythrin (PE)-conjugated sarcomeric α-actinin (SA) antibody (Miltenyi Biotec Australia Pty) or PE-conjugated mouse isotype (IgG) control (Miltenyi Biotec Australia Pty). Stained samples were analysed on a FACS CANTO II (Becton Dickinson) machine with FACSDiva software (BD Biosciences). Data were analysed was using FlowJo software and cardiac populations were determined with population gating from isotype controls.

### Immunohistochemisty/TUNEL stain

After overnight treatment at low pH, replated hiPSC-CMs were fixed in 4% paraformaldehyde for 10 min, washed in PBS, and incubated for 1 h at room temperature in blocking solution (PBS containing 2% heat-inactivated sheep serum (HISS) and 0.05% Triton X-100). Cells were incubated overnight at 4°C with primary antibody followed by incubation with secondary antibody for 1 h at room temperature. Nuclei were counterstained with 1 μg/mL DAPI. The samples were washed with PBS and TUNEL stain (Abcam) was performed as per manufacturer protocol. Stained samples were imaged within one week. All antibody dilutions were prepared in blocking solution. The following antibodies were used: mouse anti-α-actinin (Clone EA-53, Sigma Aldrich, 1:100) and donkey anti-mouse AlexaFluor 647 (ThermoFisher, 1:100). High-resolution images were obtained using a Zeiss LSM710 AiryScan confocal microscope with 20x 0.8 NA or 40x 1.3 NA Plan Apochromat objectives running Zeiss Zen Black. All image processing and quantitation was performed in FIJI^75^

### qRT-PCR

To assess mRNA transcript levels in hiPSC-CM populations, total RNA was extracted using a RNeasy Mini Kit (QIAGEN). Superscript III First Strand Synthesis (ThermoFisher) was used to generate cDNA and qRT-PCR was performed on ViiA 7 Real-Time PCR Machine (Applied Biosystems) with SYBR Green PCR Master Mix (ThermoFisher). Transcript copy numbers were calculated using the 2-ΔΔCt method relative to the housekeeping gene *HPRT1*. Primers used for qRT-PCR analysis were as follows:

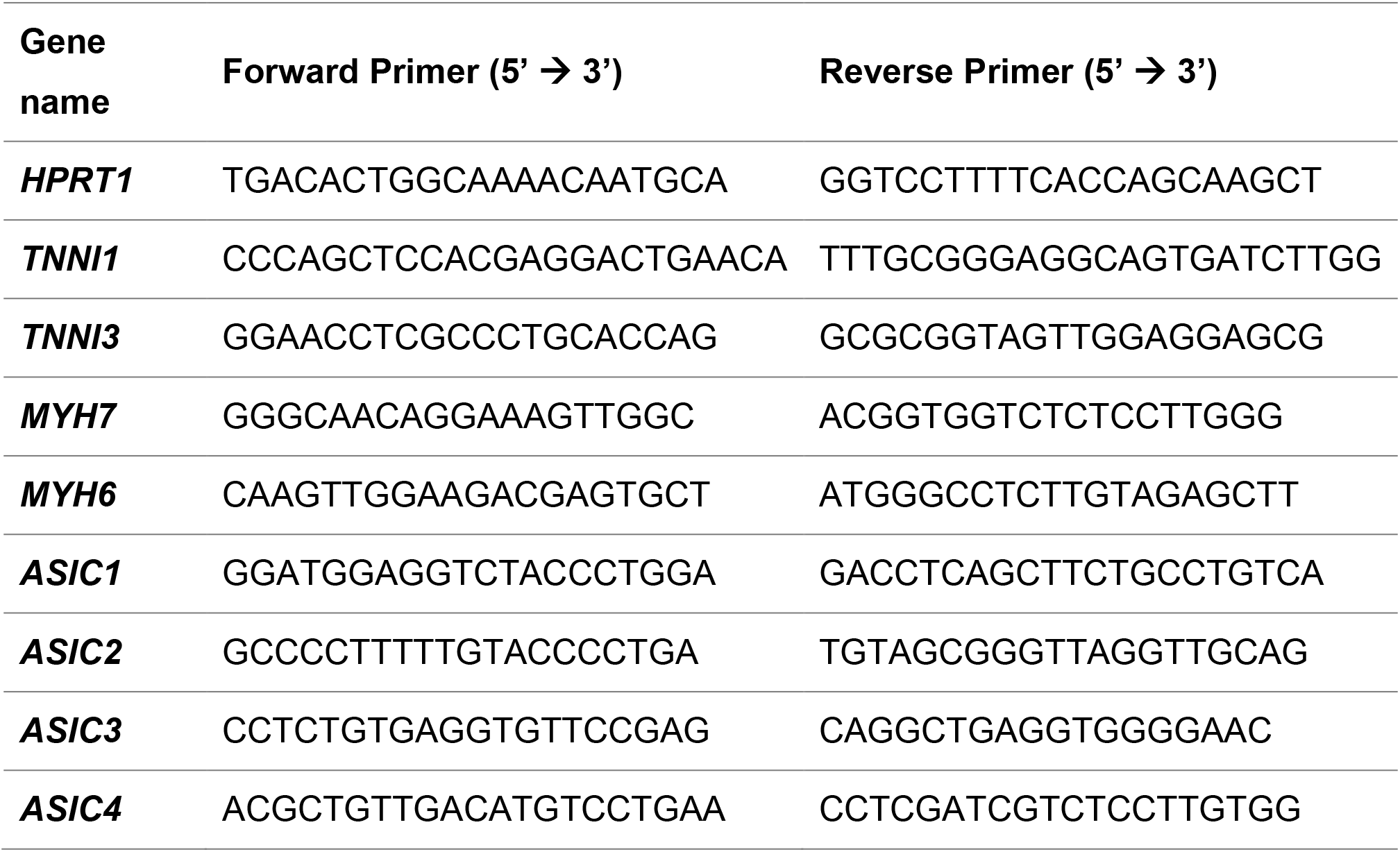

### hiPSC-CM calcium analysis

Cardiomyocytes were replated (2 x 10^4^ per well) in a 384 well plate (CellBIND black with clear bottom, Corning) and cultured for 7 days in RPMI + B27. On the day of the experiment, the cells were loaded for 1.5 h at 37°C with FLIPR calcium 4 dye (Molecular Devices) diluted in HBSS pH 7.4. After loading, the plate was transferred to a FLIPR Tetra fluorescent plate reader (Molecular Devices). Calcium transients were measured with excitation wavelengths at 470–495 nm and emission at 515–575 nm. For each plate, the camera gain and intensity were adjusted to optimize signal intensity. Data for each well were expressed as normalized arbitrary fluorescence units. All data were acquired at 0.5 s per read, with baseline measurements for 45 s followed by at least 100 s of data collection after each peptide addition. To assess peptide effects on calcium handling in normal physiological conditions, concentrated peptide dilutions (in HBSS pH 7.4) were added to each well to give a final concentration of 1 nM, 10 nM, 100 nM, or 1 μM. Calcium amplitude, maximum calcium, minimum calcium, and spontaneous beating rate were analysed using ScreenWorks software (Molecular Devices) and normalized to baseline measurements.

### Analysis of ASIC1 from GWAS studies of relevant conditions

To assess whether genetic variation of *ASIC1* associates with cardiovascular disease and stroke, we performed a gene-based level test on GWAS summary data using fastBAT^49^ implemented in the Complex-Traits Genetics Virtual Lab (CTG-VL)^76^. GWAS summary data contains the statistical information of the association of all the genetic variants included in a GWAS against a particular trait. fastBat tests the aggregated effects of a set of genetic variants within or close to (± 50 kb) each tested gene (*ACCN1* in this case) using a set-based association approach which accounts for the correlation between genetic variants (i.e. linkage disequilibrium). This provides a more powerful approach over single-variant tests done in GWAS. Specifically, we performed analyses using GWAS summary data for acute MI (N_cases_ =5,948, N_controls_ = 354,176), major coronary heart disease (N_cases_ = 10,157, N_controls_ = 351,037) and MI (N_cases_ = 7,018, N_controls_ = 354,176) from Neale’s UK Biobank GWAS database^48^ and stroke (including any type of stroke, ischemic stroke, large artery stroke, cardioembolic stroke and small vessel stroke^50^ (N_cases_ =40,585, N_controls_ = 406,111).

### Computational analysis of human translatome data

Transcriptional (mRNA-seq) and translational (Ribo-seq) raw data of human heart were obtained from the Hubner Lab website (http://shiny.mdc-berlin.de/cardiac-translatome/)^46^. Quality control checks for both RNA-seq and Ribo-seq data sets were performed to ensure reasonable quality for downstream analysis. Genes that were expressed in > 99% of samples were retained. After quality control, 47,094 and 43,183 genes were retained in the RNA-seq and Ribo-seq datasets, respectively. DESeq2 normalization was applied to normalize sequencing depth and RNA composition. The detailed normalization workflow performed by DESeq2^77^ involved four steps. First, calculated row-wise geometric mean (pseudo-reference sample) by square root of the product of all expression values across samples for each gene. Second, the ratio of each sample to the row-wise geometric mean was calculating by dividing the expression value of every gene by their row-wise geometric mean within a sample. Third, the normalization factor was calculated for each sample, which is the median value of all ratios for a sample. Lastly, the normalized count was calculated using the normalization factor, in which each raw count in a sample was divided by the normalization factor of that sample, and this was performed for every gene in every sample. The normalized counts were log-transformed to eliminate the effect of extreme values.

A total of 80 samples were analysed, including a dilated cardiomyopathy group (65 samples) and control group (15 samples). We utilized R package ggplot2 (v3.2.1)^78^ with box and violin plot to generate the two conditions in one graph for the genes encoding ASIC1, ASIC2, ASIC3, and ASIC4, in order to visualize the distribution of gene expression within one group and compare expression differences between groups.

## Notes

### Competing Interest Statement

The authors have declared no competing interest.

