## Supplementary Material for "Acid sensing ion channel 1a is a key mediator of cardiac ischemia-reperfusion injury"

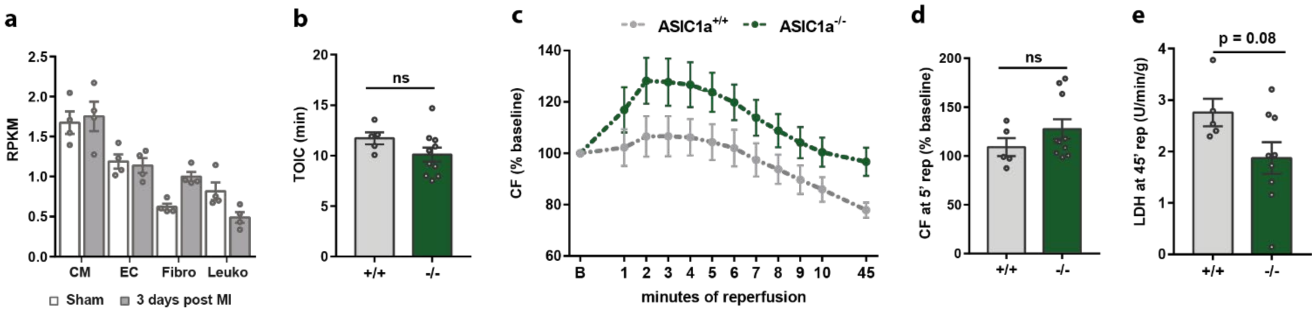


**Supplementary Fig. 1**. *Ex vivo* IRI in Langendorff-perfused ASIC1a KO mouse hearts compared to wildtype. **(a)** mRNA expression (reads per kilobase million, RPKM) analysis of ASIC1 in sorted cardiac cell populations from sham or day 3 post MI in P56 adult mouse hearts (data extracted from Ref.[1]). **(b-e)** Hearts from ASIC1a KO (ASIC1a$^{-/-}$, *n* = 10, dark gray) and WT (ASIC1a$^{+/+}$, *n* = 5, light grey) mice were subjected to 25 min of global ischemia followed by 45 min of reperfusion. **(b)** Time to onset of ischemic contracture (TOIC, p = 0.162). **(c)** coronary flow (CF) at baseline (B, pre-ischemia), during the first 10 min of reperfusion, and at the end of the 45 min reperfusion period. **(d)** CF at 5 min reperfusion (p = 0.26). **(e)** Cell death after 45 min of reperfusion (units of LDH normalized to reperfusion flow rate and heart weight, U/min/g, p = 0.08). For all parameters, baseline values were obtained immediately prior to ischemia, and all data are expressed as mean ± SEM.

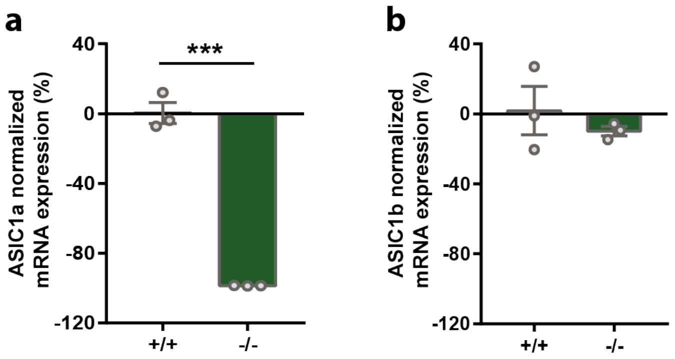


**Supplementary Fig. 2**. Verification of the loss of ASIC1a in knockout mouse strain. **(a-b)** Normalized mRNA expression levels of **(a)** ASIC1a ($p<0.0001$, WT 0.3 ± 6.0%, ASIC1a$^{-/-}$ –98.7 ± 0.7%) and **(b)** ASIC1b (p=45, WT 1.9 ± 13.8%, ASIC1a$^{-/-}$ –9.9 ± 2.6%) from brain samples from WT (black, *n* = 3) and ASIC1a$^{-/-}$ mice (green, *n* = 3). Statistical significance was determined with a two-tailed unpaired student's *t*-test (***$p<0.001$). Data are presented as mean ± SEM.

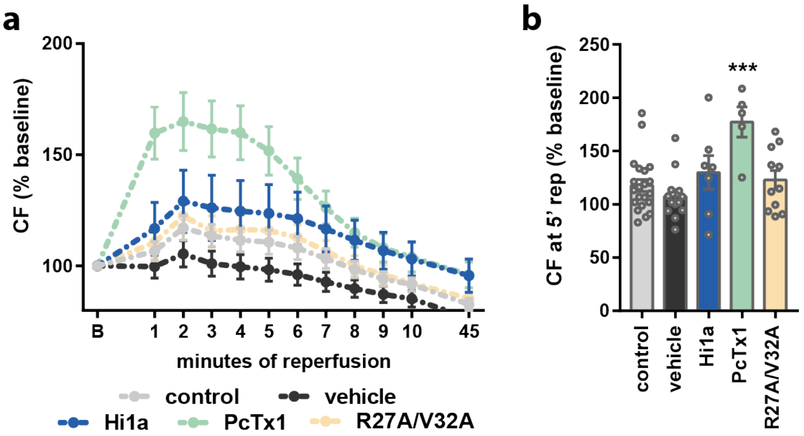


**Supplementary Fig. 3**. **ASIC1a inhibitors protect mouse hearts from *ex vivo* IRI. (a-b)** Additional analysis of the experiment described in Figure 2. **(a)** CF plotted versus time (min) at baseline (B, pre-ischemia), during the first 10 min of reperfusion, and at the end reperfusion (45 min). **(b)** CF at 5 min reperfusion. Statistical significance was determined with one-way ANOVA (***p < 0.001). Data are presented as mean ± SEM (*n* > 5/group).

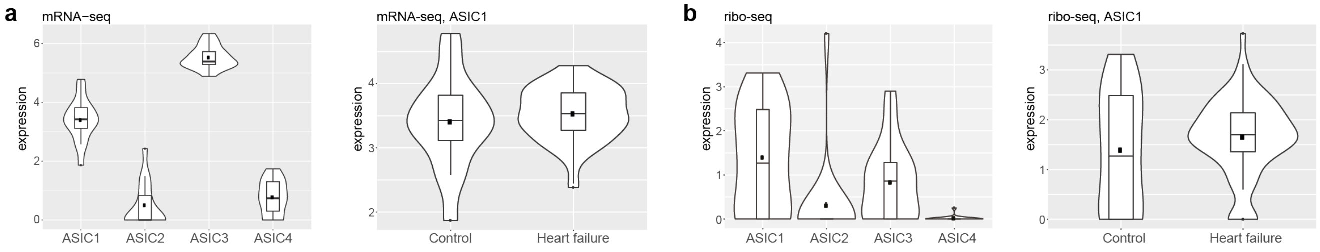


**Supplementary Fig. 4**. ASIC expression in adult human heart. Analysis of published **(a)** transcriptomic (mRNA-seq) and **(b)** translatomic (ribo-seq) data from the left ventricles of control (*n* = 15) and heart failure (dilated cardiomyopathy) (*n* = 65) patients[2].

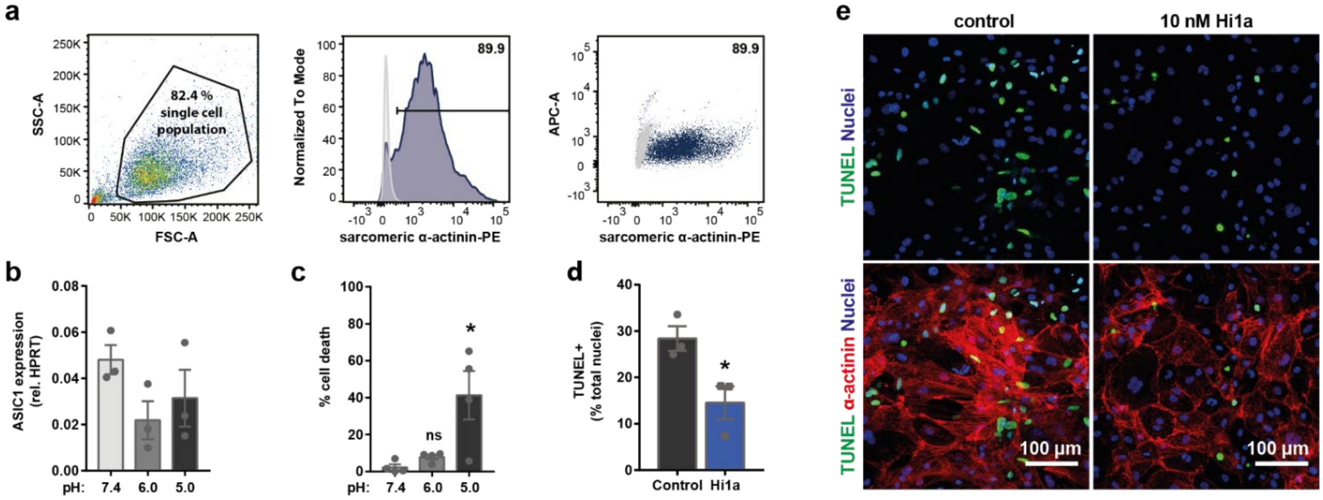


**Supplementary Fig. 5**. Generation of hiPSC-CMs and treatment at low pH. **(a)** Flow cytometry analysis of differentiated hiPSC-CMs prior to replating. Single cell population from SSC-A (side scatter) versus FSC-A (forward scatter) plot (left) was analysed for the percentage of cells that stained positive for sarcomeric α-actinin (PE-gated population) shown as a histogram (middle) and a scatter plot against an unstained fluorophore (right). Isotype-stained sample (grey) was used to create PE+ gate to analyse sarcomeric α-actinin stained sample (blue). **(b-c)** Replated hiPSC-CMs treated overnight in HBSS at pH 7.4, 6.0, or 5.0 and analysed for **(b)** mRNA expression (qRT-PCR) of ASIC1 and **(c)** cell death (LDH). **(d-e)** Replated hiPSC-CMs treated overnight in HBSS at pH 5.0 and evaluated for cell death. **(d)** Quantification of cell death (TUNEL-positive nuclei normalized to total nuclei) following **(e)** immunohistochemistry for TUNEL (green) and α-actinin (red) with nuclei counterstained with DAPI. Top panel: TUNEL and DAPI merged image. Bottom panel: TUNEL, α-actinin, and DAPI merged image. All data are expressed as mean ± SEM ($n$ = 3 biological replicates, 2–3 technical replicates each). Statistical significance was determined with one-way ANOVA (LDH results, panel c) or with a two-tailed unpaired student's *t*-test (TUNEL quantification, panel d) (*$p < 0.05$).
